# Prevalence and factors associated with Skilled Birth Attendants use among post-partum women in Arusha Tanzania

**DOI:** 10.1101/322032

**Authors:** Febronia Lawrence Shirima, Aurelia Severine Komba, Sia E Msuya, Michael Johnson Mahande

## Abstract

**Introduction:** Death related to pregnancy complications and delivery remains a public health problem in developing countries including Tanzania where majority of women and newborns die each year due to preventable causes. Delivery with a skilled birth attendant (SBA) is an important intervention to avert maternal and newborn mortality. This study aimed to determine factors influencing skilled birth attendant’s use among postpartum women in Arusha city and Arumeru district in Arusha region.

**Methods:** This was a cross-sectional study which was conducted from April to July 2017. The study utilized secondary data from the parent study which was conducted in Arusha city and Arumeru district in Arusha region. The parent study aimed to assess knowledge of danger signs in obstetric complication, birth preparedness and utilization of skilled birth attendants among post-partum women in urban and rural Arusha region in Tanzania. Data analysis was performed using SPSS version 20. Chi square statistics was used to determine the association between a set of independent variables and utilization of skilled birth attendance in bivariate analysis. Odds ratios with 95% confidence interval for factors influencing skilled birth attendants’ use were estimated in a multivariable logistic regression models. A p-value of less than 0.05 was considered statistically significant.

**Results:** A total of 1218 post-partum women were studied. Their mean age was 26.2 (SD 5.9) years. Of these, 1020 used skilled birth attendants during delivery; this corresponds to prevalence of 83.7%. Mother’s education level, area of residence, distance from health facility, frequency of antenatal care and parity were significantly associated utilization of skilled birth attendants during delivery.

**Conclusion:** The prevalence of skilled birth attendants’ use among postpartum women is high in the studied population compared to the national level. Strategies to create community awareness on important of skilled birth attendants’ use will increase its uptake and my help to reduce both mother and child mortality.

## Introduction

Globally, the maternal mortality ratio is estimated to be 261 maternal deaths per 100,000 live births with majority (99%) of these deaths occurring in developing countries especially sub Saharan Africa, where 66% of the maternal deaths occur^1^. This shows that women in the developing countries continue to lose their lives due to preventable causes of pregnancy and delivery complications. The WHO has made a call to reduce global mortality rate to less than 70 per 100,000 live births by 2030^2^. Furthermore a supplementary national target was made that no country should have a maternal mortality more than twice the global average by 2030^3^. Access to skilled birth attendants (SBA) during delivery can make the difference between life and death of mother and child hence accelerating the achievement of the global goal of 70 per 100000 live births by 2030.

Skilled birth attendants (SBA) refers exclusively to people with midwifery skills (doctors, midwives, nurses) who have been trained to proficiency in the skills necessary to manage normal deliveries and diagnose or refer obstetric complication^4^. Delivery by SBA also has impact on child survival. Deaths in the first month of life, the neonatal period account for over 44% of all deaths among children under five years of age^5^. Studies have shown that a properly trained SBA, could reduce mortality that occurs during labor, delivery and shortly after birth by providing clean delivery practices, labor surveillance, neonatal resuscitation to address intra-partum-related birth asphyxia and treatment of infections^6^

Globally, the proportion of deliveries attended by SBA increased from 59% to 71%, yet this leaves more than one in four babies and mothers without access to this crucial medical care during child birth^7^. As a result, maternal and newborn survival rate still remains low. There exists a variation in SBA coverage, it is much lower (52%) in southern Asia and sub-Saharan

Africa^7^. This variation makes the maternal morbidity and mortality rate to remain comparatively high in developing countries including Tanzania. The current SBA coverage in Tanzania is 64%^8^. This is slightly lower than the global average.

In Tanzania under the national road map strategic plan to improve reproductive, maternal, new born, child and adolescent health, the ministry of health has set target to increase coverage of deliveries attended by skilled birth attends from 51% to 80% by 2020 by putting more emphasis in the provision of quality health services offered by skilled attendants in enabling environment^9^. In order to achieve this target, more emphasis should be done to identify and eradicate the barriers which make expectant mothers not to access SBAs or make them to prefer home delivery. The aim of this study was to determine factors affecting SBA use among postpartum women in Arusha city and Arumeru districts.

## Materials and Methods

### Study design and data source

This was a cross-sectional study which was conducted from April to July 2017. The study utilized secondary data from the parent study which was conducted in Arusha city and Arumeru district in Arusha region which aimed at assessing knowledge of danger signs in obstetric complication, birth preparedness and utilization of skilled birth attendants among post-partum women in urban and rural Arusha region in Tanzania.

### Description of the parent study

The purpose of the parent study was to assess knowledge of danger signs in obstetric complication, birth preparedness and utilization of skilled birth attendants among post-partum women in urban and rural Arusha region in northern Tanzania. The study addressed more about knowledge of danger signs in obstetric complications and birth preparedness, factors associated with having birth preparedness and complication readiness plan among post-partum women. The study also looked into prevalence of women with knowledge of danger signs and prevalence of women who had birth preparedness plan. However the study did not address about factors influencing SBA use among post-partum women. Therefore the current study aimed at determining factors influencing SBA use.

### Study area

The parent study was carried out in two districts; Arusha city and Arumeru district of Arusha region. The region has a total of 696,320 people (census report, 2013). The two districts have a total of 110 health facilities. Arusha city has 3 hospitals, 5 health centers and 45 dispensaries making a total of 58 health facilities. Arumeru district has 2 hospitals, 5 health centers and 45 dispensaries making a total of 52 health facilities.

### Study population

Post-partum women attending Reproductive Child Health (RCH) services in selected health facilities were studied. The study included post-partum women with infants aged less than 12 months of age attending selected health facilities for routine post natal care in the selected health facilities and those who consented to participate in the study. The study excluded post-partum women who did not consent to participate in the study and those who were not permanent residents in the two districts. Women who did not give information concerning SBA use were also excluded

### Sample size and sampling technique

The study enrolled a total of 1251 participants. Two districts were randomly selected out of six districts of Arusha region. Selection of facilities in the two districts was based on large number of clients attending the RCH clinics. Eight health facilities with largest number of post natal care attendants in each district were selected. At the clinics all women meeting inclusion criteria were invited to participate.

### Data collection methods and tools

The women attending RCH clinic were interviewed. A questionnaire was used which consisted of both closed and open ended questions. Information collected included socio-demographic, economic variables, reproductive history, number of antenatal care (ANC) visits, and information on delivery, as well as assessment level of knowledge on modern contraceptives during post-natal period.

### Study variables

The skilled birth attendant use during delivery was the main outcome in this study. The independent variables includes; socio-demographic variables (age, education, residency, marital status, income, occupation), maternal factors (gravidity, parity) and healthcare related factors (ANC attendance, frequency of ANC attendance)

### Data analysis

Data analysis was performed using SPSS version 20. Descriptive statistics was summarized using frequency and proportions for categorical variables. Chi square statistics was used to determine the association between a set of independent variables and utilization of skilled birth attendance in bivariate analysis. Odds ratios (ORs) with 95% confidence interval (CI) for the factors influencing skilled birth attendant use was estimated in a multivariable logistic regression models. A p-value of less than 5% (2-sided) was considered statically significant.

### Ethical consideration

The ethical approval was obtained from Kilimanjaro Christian Medical University College research and ethics committee. Permission for using the parent study data was obtained from the institute of public health of Kilimanjaro Christian Medical University College. Confidentiality was adhered where women identification number were used instead of names.

## Results

### Characteristics of study participants

The general characteristics of study participants are shown in table 1. A total of 1218 postpartum women were studied, half of them 625 (51.3%) were from Arusha municipal and the rest were from Arumeru district. The mean age of the participants was 26.2 (SD 5.9) years. Majority (71%) lived less than an hour away from a health facility. More than half of them (56.2%) had primary education and majority (79.5%) of the women were either married or cohabiting. Most of the participants were unemployed (90.8%). Majority (68%) of the participants earned an income of <60000 Tanzanian shillings per month. Most (85.5%) women had less than four alive children and gravidity ranged from 0–3 (81.4%). Most (99.6%) of the women had attended ANC during their last pregnancy and more than a half (67.7%) had attended ANC more than four times.

**Table 1:** Characteristics of study participants. (N=1218)

| Characteristic | n | % |
| --- | --- | --- |
| Age(in years)* |  |  |
| <20 | 134 | 11.0 |
| 20-34 | 955 | 78.4 |
| >34 | 128 | 10.5 |
| Education level |  |  |
| Never been to school | 117 | 9.6 |
| Primary | 684 | 56.2 |
| Secondary | 360 | 29.6 |
| Tertiary | 57 | 4.7 |
| Marital status |  |  |
| Married/ cohabiting | 1143 | 93.8 |
| Single/divorced/widow | 85 | 6.9 |
| Residence |  |  |
| Urban | 625 | 51.3 |
| Rural | 593 | 48.7 |
| Employment on monthly payment |  |  |
| Employed | 112 | 9.2 |
| Non employed | 1106 | 90.8 |
| Personal income (in shillings) |  |  |
| ≤60000 | 828 | 68.0 |
| 61000-200000 | 271 | 22.2 |
| 201000-500000 | 71 | 5.8 |
| >500000 | 48 | 3.9 |
| Distance to health facility |  |  |
| <60min | 865 | 71 |
| 61-180min | 288 | 23.6 |
| >181min | 65 | 5.3 |
| ANC attendance |  |  |
| No | 5 | 0.4 |
| Yes | 1213 | 99.6 |
| ANC visits |  |  |
| < 4 | 394 | 32.3 |
| ≥4 | 824 | 67.7 |
| Parity |  |  |
| Less than 4 | 1041 | 85.5 |
| 4 and above | 177 | 14.5 |
| Gravidity |  |  |
| 0-3 | 991 | 81.4 |
| 4 and above | 227 | 18.6 |

\*1 missing

### Prevalence of skilled birth attendant use

Among the 1218 women, 1020 used skilled birth attendants during their last delivery. This corresponds to the prevalence of SBA use of 83.7%.

### Bivariate analysis for factors associated with SBA use

The bivariate analyses for factors associated with SBA are shown in table 2. Numerous social demographic factors were significantly influencing SBA use. These include area of residence (p< 0.001), age (p=0.036), maternal education level (p<0.001), marital status (p=0.008), employment (p<0.001) and distance to the nearest facility (p=<0.001). In additional, some obstetrics characteristics such as gravidity (p<0.001), parity (p<0.001) and healthcare related factors such as ANC attendance (p<0.001), ANC frequency (p<0.001) were found to be significantly associated with SBA use.

**Table 2:** Factors affecting SBA use; bivariate analysis (N=1218)

| Variable | SBA<br>(N) | USE<br>n(%) | P-value |
| --- | --- | --- | --- |
| **Residency** |  |  |  |
| Urban | 625 | 604(96.6) | <0.001 |
| Rural | 593 | 416(70.2) |  |
| **Age** |  |  |  |
| <20 | 134 | 105(78.4) | 0.036 |
| 20-34 | 955 | 814(85.2) |  |
| >35 | 128 | 101(78.9) |  |
| **Marital status** |  |  |  |
| Married/cohabiting | 1143 | 949(83.0) | 0.008 |
| Single/divorced/widow | 75 | 71(94.7) |  |
| **Education** |  |  |  |
| Never been to school | 117 | 42(35.9) | <0.001 |
| Primary | 684 | 576(84.2) |  |
| Secondary and tertiary | 417 | 402(96.4) |  |
| **Employment** |  |  |  |
| No | 1106 | 910(82.3) | <0.001 |
| Yes | 112 | 110(98.2) |  |
| **distance to the nearest facility** |  |  |  |
| <60min | 865 | 746(86.2) | <0.001 |
| 60-180min | 288 | 230(67.7) |  |
| >180min | 65 | 44(67.7) |  |
| **ANC attendance** |  |  |  |
| no | 5 | 4(80.2) | 0.589 |
| yes | 1213 | 1016(83.8) |  |
| **ANC frequency** |  |  |  |
| >4 | 394 | 289( 73.4) | <0.001 |
| 4 and above | 824 | 731(88.7) |  |
| **Parity** |  |  |  |
| >4 | 1041 | 906(87.0) | <0.001 |
| 4 and above | 177 | 114(64.4) |  |
| **Gravidity** |  |  |  |
| 0-3 | 991 | 862(87.0) | <0.001 |
| 4 above | 227 | 158(69.6) |  |

### Multivariable logistic regression analysis for factors associated with SBA use

In the multivariable logistic regression analysis (table 3), some of socio-demographic, obstetrics and health care related factors were independently associated with SBA use. Women who reported living in urban had 7 fold (OR 7.16; 95% CI [4.32–35.72]) higher odds of SBA use as compared to those living in the rural area. Women with primary education (OR 5.74; 95% CI [3.64–9.07]) and those with secondary education and above (OR 18.27; 95%CI [9.34–35.72]) were also more likely to use SBA during delivery as compared to women who had never been to school. Additionally, nearest distance to the health facility increased women’s likelihood of SBA use (OR 2.49; 95%CI [1.26–4.95]) than women who reported walking for more than 180 minutes to the health facility. In the other hand; frequency of ANC visits especially of >4 visit was associated with SBA use (OR 2.66; 95% CI [1.94–3.66]) compared with ANC of less than 4 visit.

**Table 3:** Factors influencing SBA use; multivariable analysis (N=1218)

| variable | SBA<br>N | USE<br>n(%) | COR(95%CI) | AOR(95%CI) |
| --- | --- | --- | --- | --- |
| **Place of residency** |  |  |  |  |
| urban | 625 | 604(96.6) | 12.2(7.66-19.57) | 7.16(4.32-35.72) |
| rural | 593 | 416(70.2) | 1 | 1.0 |
| **Age of participants** |  |  |  |  |
| <20 | 134 | 105(78.4) | 1.54(0.97-1.75) | - |
| 20-34 | 955 | 814(85.2) | 0.97(0.54-1.75) | - |
| >34 | 128 | 101(78.9) | 1 |  |
| **Marital status** |  |  |  |  |
| Married/cohabiting | 1143 | 949(83.0) | 0.28(0.09-0.76) | - |
| Single/divorced/widow | 75 | 71(94.7) | 1 |  |
| **Education of participants** |  |  |  |  |
| Never been to school | 117 | 42(35.9) | 1 | 1.0 |
| Primary school | 684 | 576(84.2) | 9.5(6.2-14.6) | 5.74(3.64-9.07) |
| Secondary and above | 417 | 402(96.4) | 47.8(25.1-90.7) | 18.27(9.34-35.72) |
| **Employment** |  |  |  |  |
| no | 1106 | 910(82.3) | 0.08(0.02-0.35) | - |
| yes | 112 | 110(98.2) | 1 |  |
| **distance to the nearest facility** |  |  |  |  |
| <60min | 865 | 746(86.2) | 3.0(1.72-5.21) | 2.49(1.26-4.95) |
| 60-180min | 288 | 230(67.7) | 1.9(1.04-3.43) | 1.53(0.74-3.19) |
| >180min | 65 | 44(67.7) | 1 | 1.0 |
| **ANC attendance** |  |  |  |  |
| No | 5 | 4(80.2) | 0.78(0.08-6.98) | - |
| Yes | 1213 | 1016(83.8) | 1 |  |
| **ANC frequency** |  |  |  |  |
| <4 times | 394 | 289(73.4) | 1 | 1.0 |
| 4 and above | 824 | 731(88.7) | 2.86(2.09-3.89) | 2.66(1.94-3.66) |
| **Parity** |  |  |  |  |
| <4 | 1041 | 906(87.0) | 3.7(2.59-5.29) | 3.25(1.45-7.31) |
| 4 and above | 177 | 114(64.4) | 1 | 1.0 |
| **Gravidity** |  |  |  |  |
| 0-3 | 991 | 862(87.0) | 2.9(2.08-4.09) | - |
| 4 and above | 227 | 158(69.6) |  |  |

Furthermore, women who reported having less than four children had 3 folds (OR 3.25; 95% CI [1.45–7.31]) higher odds of SBA use as compared their counterparts who had four or more children. Other factors such as maternal age, marital status, employment status, attendance to ANC and gravidity were not associated with SBA use after adjusting for other factors in the models

### Discussion

SBA use was high in the study area, with prevalence of 83.7%. This proportion is higher than the national estimate of 64%^8^. It is also higher than other studies which have been done in Tanzania^10^ and other studies in Uganda^11^, in Ghana^12^ and Ethiopia^13^. The reasons given for not using SBA were; being forced by family members, being denied by husband, self-made decision, transport problem, bad weather, hospital being far away and sudden onset of labor. These reasons are avoidable if education concerning the importance of SBA is provided to the family members and the community at large. This also calls for the government to take part in ensuring good infrastructures are built in remote areas.

This study has found education status to be significantly associated with SBA utilization such that, women with higher education have higher tendency of SBA use. Similar findings were previously reported in Tanzania^14,15^ and India^16^. These findings show that attainment of higher education, helps women to become more aware of pregnancy associated risks and consequences; hence they tend to seek proper health care through SBA during delivery as compared with uneducated women who may have limited access to information.

Distance from the health facility was associated with skilled birth attendant use during delivery. This finding is similar to a study done in Nepal which showed that women who walk for more than an hour to a health facility with delivery services are less likely to use SBA during delivery^17^. This is also similar to a study done in Uganda^11^. Living less than an hour away from the hospital guarantees short time taken by the mother to reach the facility when labour starts and a lower transport cost. This motivates facility delivery under a skilled birth attendant.

In the present study ANC frequency was associated with SBA use during delivery such that women who had more than 4 visits were more likely to deliver with SBA. This finding is consistence with previously reported by the study in Nepal^19^. Significance of ANC frequency in SBA utilization suggests that if ANC is sought properly as per WHO recommendation, may result to awareness creation of the danger signs and even the services received encourage women to seek appropriate delivery care.

Parity of the mother was significantly associated with SBA use during delivery. Multiparous women were less likely to use SBA during delivery. This finding is similar to previous studies in Tanzania^16^ and Ethiopia^20^. This study is also in agreement with a study done in Uganda^21^. This shows that multiparous women are more confident on child birth, compared to prime gravida women, thus they are associated with low SBA use.

### Study strengths and limitations of the study

This study used a large sample size; hence high precision and the results can be generalized to the whole Arusha region. This was a secondary data analysis; which saved the time and it was cost-efficient as data was already available.

Data was collected from women attending at the health facility, these are most likely the women who have proper health care seeking behavior, and hence they may overestimate the prevalence of SBA use.

## Conclusions

Prevalence of SBA use during delivery in Arusha city and Arumeru district is high above the national estimate of 64%. The factors associated with SBA use are:-education status, residency, distance from the health facility, ANC frequency and parity.

## Author’s contribution

FLS: Designed the study, performed statistical analysis, participated in writing the manuscript

ASK: Contributed in drafting of the manuscript, MJM: contributed in designed the study, reviewed the manuscript for intellectual contents.

## Acknowledgement

We gratefully acknowledge the department of epidemiology and biostatistics, institute of public health, Kilimanjaro Christian Medical University Collage for allowing us to use their data.

